# *7-UP:* generating *in silico* CODEX from a small set of immunofluorescence markers

**DOI:** 10.1101/2022.06.03.494624

**Authors:** Eric Wu, Alexandro E. Trevino, Zhenqin Wu, Kyle Swanson, Honesty J. Kim, H. Blaize D’Angio, Ryan Preska, Gregory W. Charville, Piero D. Dalerba, Umamaheswar Duvvuri, Jelena Levi, A. Dimitrios Colevas, Nikita Bedi, Serena Chang, John B. Sunwoo, Aaron T. Mayer, James Zou

## Abstract

Multiplex immunofluorescence (mIF) assays multiple protein biomarkers on a single tissue section. Recently, high-plex CODEX (co-detection by indexing) systems enable simultaneous imaging of 40+ protein biomarkers, unlocking more detailed molecular phenotyping, leading to richer insights into cellular interactions and disease. However, high-plex imaging can be slower and more costly to collect, limiting its applications, especially in clinical settings. We propose a machine learning framework, *7-UP,* that can computationally generate *in silico* 40-plex CODEX at single-cell resolution from a standard 7-plex mIF panel by leveraging cellular morphology. We demonstrate the usefulness of the imputed biomarkers in accurately classifying cell types and predicting patient survival outcomes. Furthermore, *7-UP’s* imputations generalize well across samples from different clinical sites and cancer types. *7-UP* opens the possibility of *in silico* CODEX, making insights from high-plex mIF more widely available.

## Introduction

The tissue microenvironment (TME) is a complex milieu comprising many cell types and heterogeneous cell states. Common techniques for understanding the TME like mass spectrometry^1^ and flow cytometry^2^ allow for bulk measurements of many cell biomarkers but discard valuable spatial information in the process. Recently, multiplexed molecular imaging assays have enabled the quantification of cell types and molecules in their native tissue context. Commercial multiplexed immunofluorescence (mIF) systems are increasingly commonplace in clinical diagnostic and prognostic settings^3^ but are typically limited to quantifying between 1 and 7 biomarkers^4^.

More recently, mIF techniques such as co-detection by indexing (CODEX)^5^ quantify 40 or more markers in situ, allowing a richer and more holistic characterization of the TME and its underlying cell types and disease processes. However, CODEX systems are significantly more costly and time-consuming to run when compared to most low-plex systems, which limits their wider adoption in clinical settings.

To address this limitation, we introduce *7-UP,* a machine learning framework that generates *in silico* high-plex mIF (30+ biomarkers) from only a panel of seven experimentally measured biomarkers. Whereas typical 7-plex measurements can only resolve up to 5-7 distinct cell types^3^, using the imputed markers from *7-UP* enable the identification of up to 16 cell types. Moreover, the imputed biomarker expressions can predict complex clinical outcomes with accuracy comparable to using experimental measurements from CODEX. *7-UP* generalizes to new cancer types and samples that come from different clinical sites than its training data. Our approach highlights a significant opportunity to use machine learning toward inferring high-dimensional molecular features from commonly available low-plex imaging data.

Imputation techniques have been applied to missing data in genomics^6–8^ and transcriptomics^9,10^ datasets, as well as in mass spectrometry and shotgun proteomics^6, 11, 12^ data. Deep learning has been used to extract morphological and spatial features from pathology H&E-stained slides^13–16^, and in turn, enabled *in silico* IHC staining^17^ and spatial transcriptomics^18^. More recently, computational methods have been developed for improving cell-type classification in CODEX-acquired data^19^ and augmenting with spatial information in particular^20^. To date, our work is the first to demonstrate the effectiveness of deep-learning-based morphological feature extraction toward clinically meaningful multiplex immunofluorescence imputation.

## Results

### 7-UP summary

The *7-UP* framework consists of the following pipeline. We first select an optimal panel of 7 biomarkers from the full CODEX biomarker panel. While the choice of which biomarkers to measure in a 7-plex imaging workflow can depend on clinician preference and disease subtype, we use a previously validated approach, Concrete autoencoder^22^, for automatically selecting informative biomarkers. This approach identified *DAPI, CD45RA, CD15, pan-cytokeratin (PanCK), HLA-DR, Ki67,* and *Vimentin* (“Main panel” in Figure 1), which we use in our main experiments. We additionally report results using an alternative panel commonly used in immunology^4,23^ consisting of *DAPI, CD4, CD15, PanCK, CD8, Ki67,* and *Vimentin* ( “Alternative panel” in Figure 1), and the results are consistent with the main panel. Next, we extract cell-level spatial features across each of these seven biomarkers in the CODEX dataset. To do this, we train a convolutional neural network^24^ to learn spatial and morphological features from cell image patches generated from the full samples. We combine cell-level spatial features with average biomarker expression values to train a machine learning regression model^25^ to impute the expression of the 30+ additional biomarkers.

**Figure 1:**
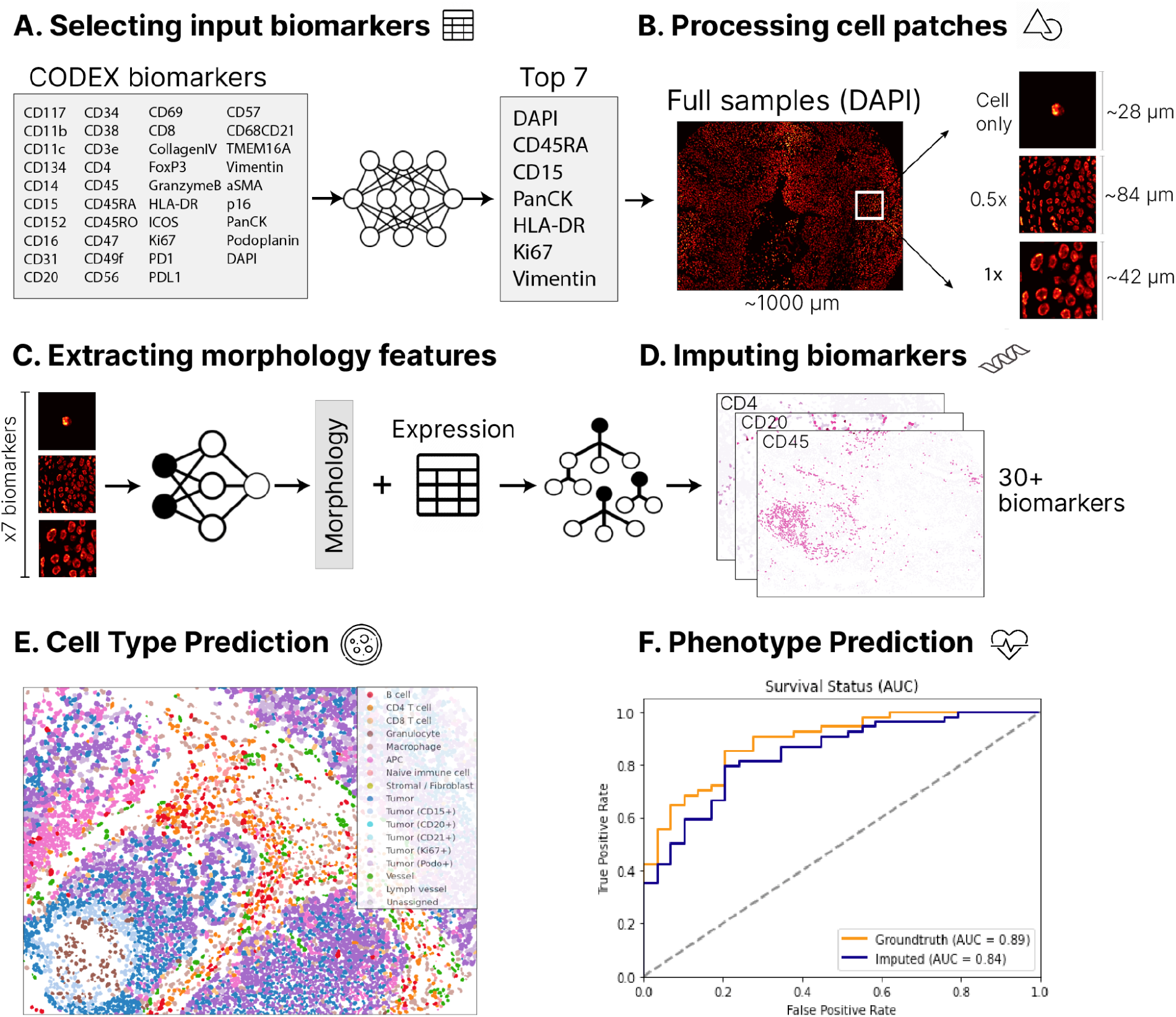
Overview of the *7-UP* Framework. **Panel A**: From the full CODEX panel of biomarkers, a Concrete autoencoder selects a subset of 7 biomarkers to reconstruct the full panel. **Panel B**: From a full sample (~1000 microns wide), image patches are extracted for each cell: *Cell only,* containing the morphology of the cell at 3x scaling, *0.5x,* a ~84-micron neighborhood around the cell, and *1x*, a ~42-micron neighborhood around the cell. **Panel C&D:** Each cell has three patches produced for each of the 7 biomarkers, totaling 21 patches used as input to a deep learning model. This model extracts morphological features for each cell, which are combined with the average expressions of the top 7 biomarkers to predict the average expressions of the remaining CODEX panel biomarkers using a machine learning regression model. **Panel E**: The imputed biomarker expressions (from Panel C&D) are used in place of the CODEX-generated values in the k-nearest neighbors algorithm used to produce cell-type ground truth. An example predicted sample is shown. **Panel F:**Using a deep learning model trained to predict phenotypic outcomes (Zheng et. al), the predicted cell types are used in place of the ground truth cell types to produce sample-level predictions for survival status, HPV status, and recurrence.

To validate the veracity of the 7-UP imputed expressions, we use them to predict cell types and patient outcomes. We replace CODEX-measured expressions with the 7-UP imputed expressions in a k-nearest neighbors algorithm used to determine cell type ground truth to generate cell type predictions. In turn, these predicted cell types are used as input in place of the CODEX-measured ground truth cell types in a graph neural network^26^ trained to produce sample-level predictions for patient-level survival status, HPV status, and recurrence.

### Application of 7-UP to head-and-neck and colorectal cancer datasets

Our primary dataset consists of 308 samples from 81 patients with head and neck squamous cell carcinomas at the University of Pittsburgh Medical Center (UPMC-HNC). Two external validation datasets are used: a head and neck squamous cell carcinomas dataset with 38 samples from 11 patients from Stanford University (Stanford-HNC) to demonstrate generalization on the same disease, and a colorectal cancer dataset with 292 samples from 161 patients from Stanford University (Stanford-CRC) to demonstrate generalization to another disease. The number of samples, patients, coverslips, and total cells in each dataset is described in Table 2. UPMC-HNC is chosen as the primary training and evaluation dataset as it contains the largest number of samples, coverslips, and total cells. We evaluate our models on held-out coverslips not seen during training to assess model robustness to technical artifacts across coverslips.

### Concordance of biomarker imputation

*7-UP* achieves an average Pearson correlation coefficient (PCC) of 0.534 across all predicted biomarkers in the UPMC-HNC dataset (Table 1). The predictive performance also holds across an alternative input panel (PCC of 0.529). Immune-related biomarkers like CD4, CD20, and CD45 are most accurately predicted, with PCCs above 0.70 (examples shown in Figure 2a).

**Figure 2:**
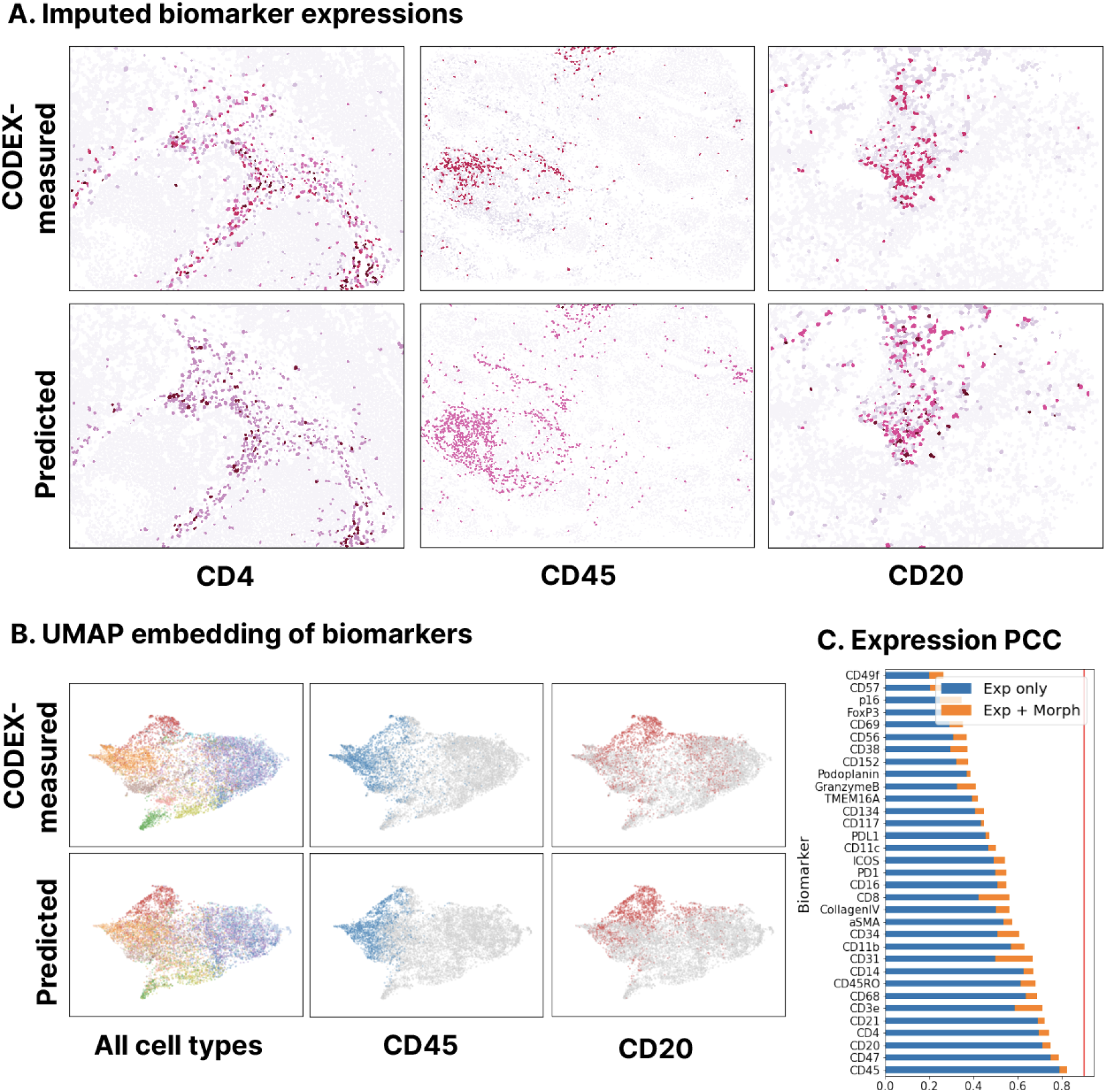
Biomarker imputation concordance on the UPMC-HNC dataset. **Panel A**: CODEX-measured versus predicted expressions for three biomarkers: CD4, CD20, and CD45. Samples shown have average patchwise PCC (Pearson correlation coefficient) scores around the 50th percentile of all samples. **Panel B:** A UMAP embedding was performed on the biomarkers of an equal sample of CODEX-measured and predicted cells. The first column is colored by the ground truth cell types (legend from Figure 1d); the second and third columns represent cells that express CD45 and CD20 (colored by expressions greater than the 75th percentile CODEX-measured value). **Panel C:** Patchwise PCC across all test samples for each biomarker. The blue bars represent the performance of a model trained only using average expression values as input; the orange bars represent the performance of a model trained using both average expression values and morphology features.

**Figure 3:**
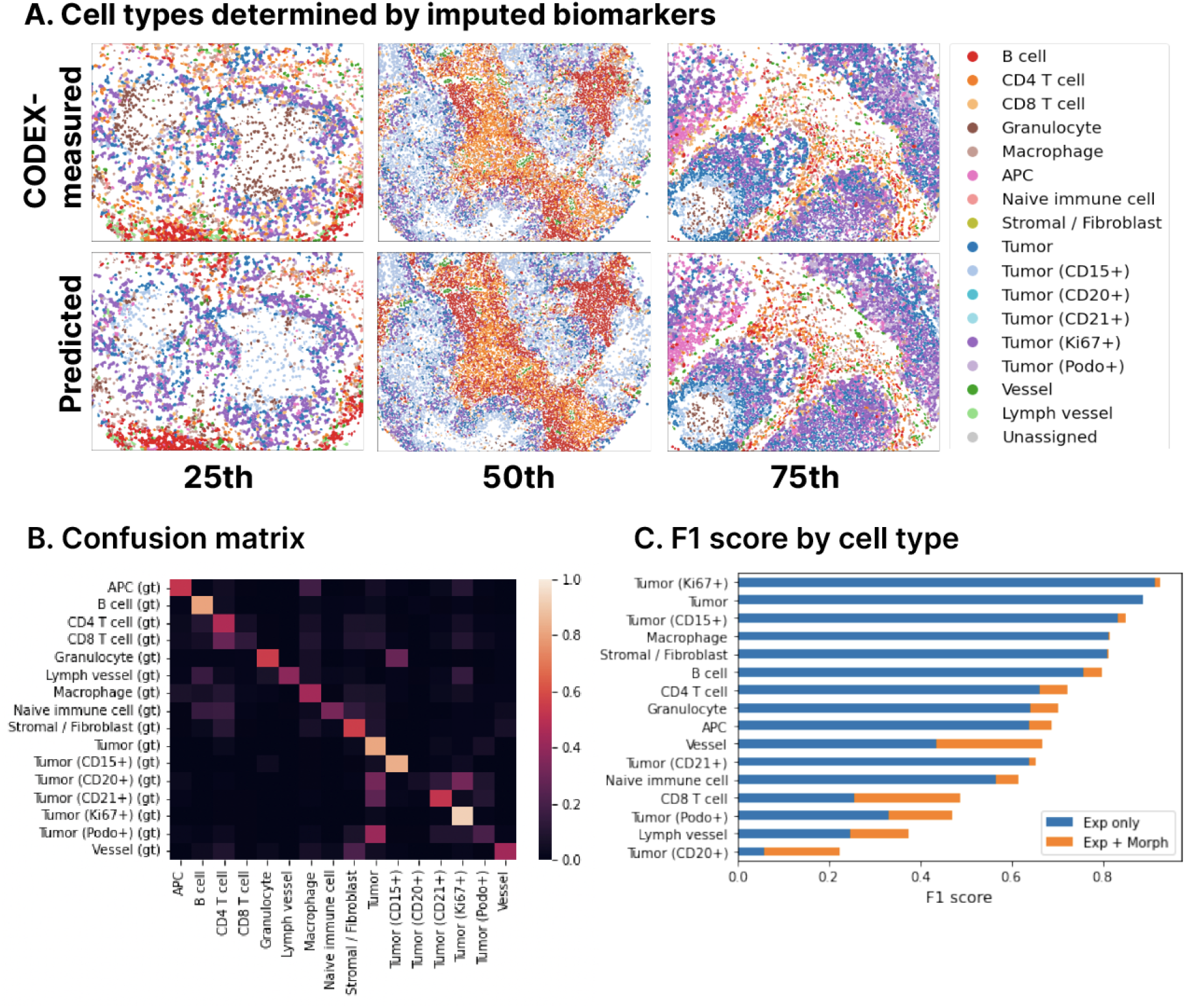
Cell type predictions closely match CODEX measurements (UPMC-HNC dataset). **Panel A:** CODEX-measured and predicted cell types are shown side-by-side on 25th, 50th, and 75th percentile samples (by patchwise F1 score). **Panel B:** Left: Confusion matrix between the kNN-determined ground truth cell types (rows) and ML imputed cell type (columns). Right: Breakdown of patchwise F1 score by cell type. The blue bars represent the performance of a model trained using only average expression values, and the orange bars represent the performance of a model trained using both average expression values and morphological features.

**Table 1:** Performance of 7-UP on the UPMC-HNC dataset. Biomarker imputation results are reported using the average patchwise Pearson correlation coefficient (PCC). Cell type predictions are reported using the patchwise weighted F1 score. The first row refers to the model trained without including morphological features in the input. The second and third rows refer to the models trained with morphological features. The main and alternative panels are described in the Methods section. Numbers in parentheses indicate the 95% bootstrapped confidence intervals.

|  | Patchwise PCC | Patchwise F1 |
| --- | --- | --- |
| **UPMC-HNC Dataset** | **33 biomarkers** | **16 cell types** |
| 7-UP model (7 biomarkers) | 0.474 (0.006) | 0.667 (0.002) |
| 7-UP model (7 biomarkers + Morphology)<br>Main panel | 0.534 (0.009) | 0.727 (0.002) |
| 7-UP model (7 biomarkers + Morphology)<br>Alternative panel | 0.529 (0.007) | 0.739 (0.002) |

### Predicting cell types from imputed biomarkers

We also measure the reliability of the imputed biomarkers by using them for determining cell types since cell type identification is a common task in analyses of CODEX data. Toward this task, *7-UP* achieves an F1 score of 0.727. The full CODEX-measured biomarker panel defines the ground truth labels in both models.

We examine how accurately the predicted cell types retain local cell neighborhood structures by comparing the spatial adjacency matrices (Supp. Figure 2). These were produced by projecting the cells into a graph representation described in Zheng et al.^26^ and then counting the relative frequencies of spatially adjacent cells. Comparing the two matrices shows that local clusters of cell types are well preserved (Root-mean-square distance of 0.0357). We additionally verify that the predicted cell types closely match the true distribution by projecting the predicted and CODEX-measured biomarker expressions using UMAP and visualizing the cell type labels (Figure 2b).

### Predicting patient phenotypes from predicted cell types

To validate the reliability of the cell types determined by *7-UP* imputed biomarkers, we use them to predict three patient phenotypic outcomes: HPV infection status, primary outcome (survival), and recurrence of disease. To this end, we use a graph-based deep learning model^26^ trained using ground truth cell types from the UPMC-HNC dataset to predict these three binary outcomes. To evaluate the veracity of our predicted cell types, we replace the CODEX-measured cell types used to make the baseline prediction with the predicted cell types as input to the model. The results demonstrate that the imputed cell types can predict phenotypic outcomes at a level comparable to the ground truth labels (Figure 4).

**Figure 4:**
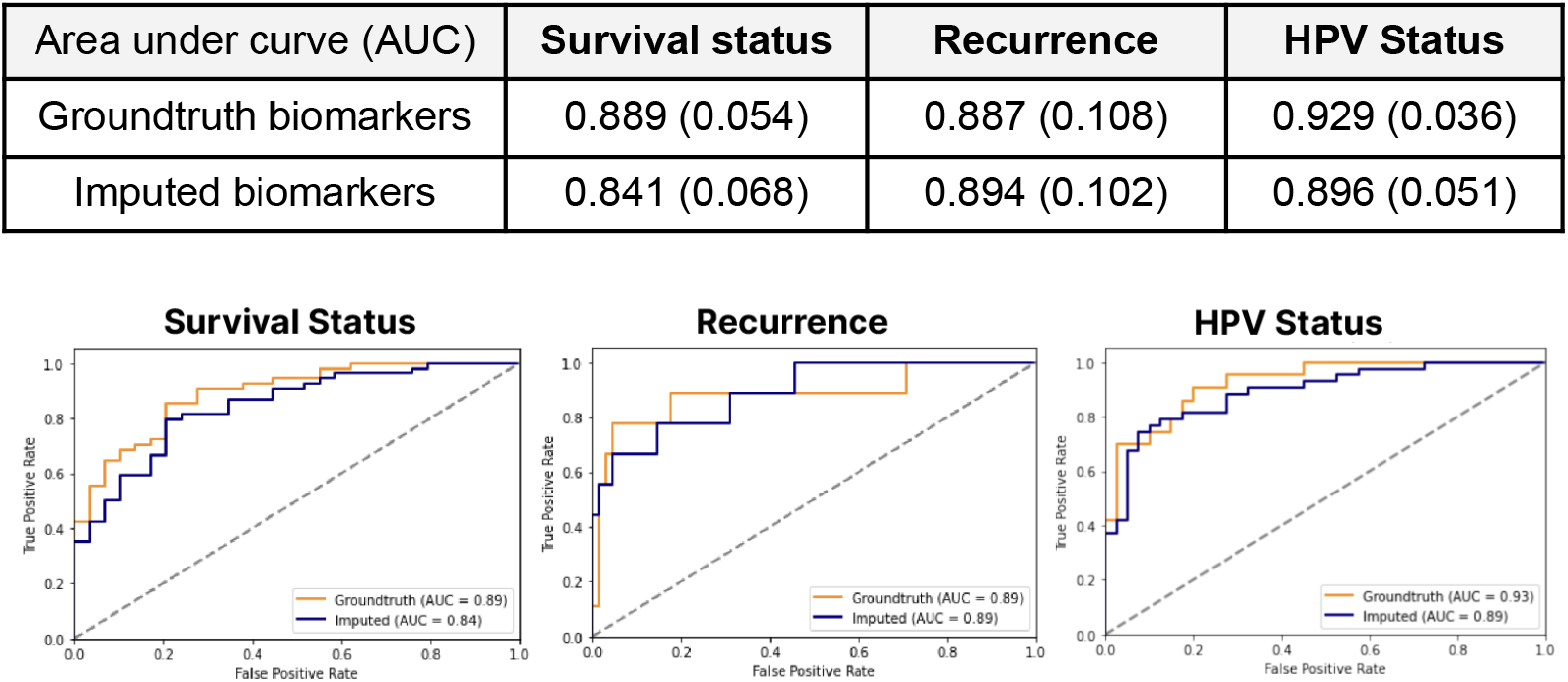
Imputed biomarkers are useful for predicting patient phenotypes (UPMC-HNC dataset). Top: Three phenotypic outcomes using imputed vs CODEX-measured biomarkers. AUC score reported (95% bootstrapped confidence interval reported in parentheses). Bottom: ROC curves of three phenotypic outcomes.

### Cross-site and cross-disease generalization

Finally, we evaluate our model on another head and neck cancer dataset (Stanford-HNC) and a colorectal cancer dataset (Stanford-CRC). The biomarker imputation and cell type prediction performances remain stable (e.g. for Stanford-CRC: 0.489 vs 0.583 PCC and 0.614 vs 0.605 F1) even when evaluated on a different clinical site and cancer type (Figure 5), indicating that the model’s performance is robust when evaluated on unseen data.

**Figure 5:**
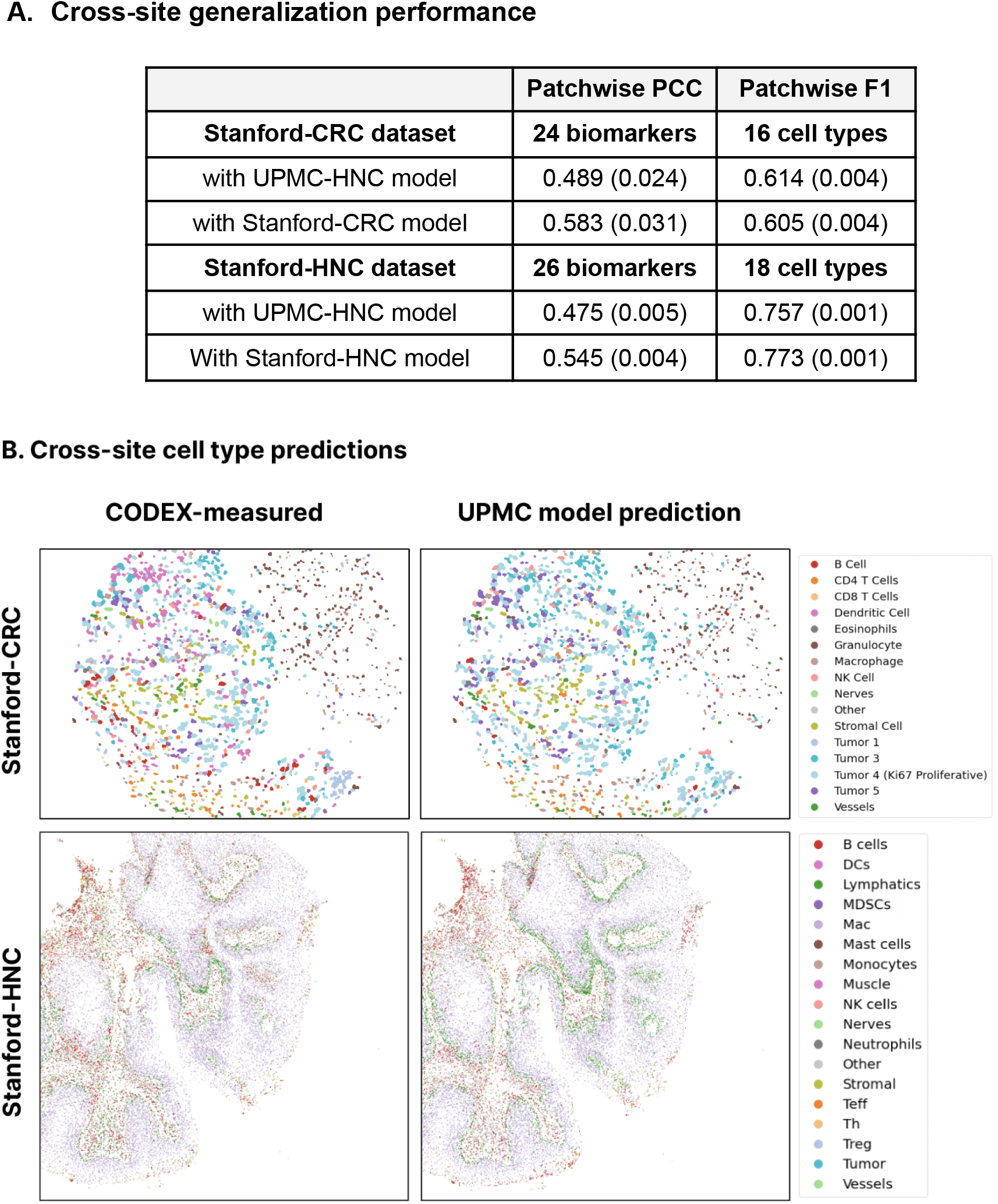
7-UP generalizes well to other data sites and disease types. **Panel A**: Imputed biomarker and predicted cell type performance on two external validation datasets (Stanford-CRC and Stanford-HNC). The UPMC-trained model’s performance is reported on each dataset, along with a reference model that has been trained on the validation dataset. Metrics are reported with 95% confidence intervals in parentheses. **Panel B:** 50th percentile (by F1 score) samples are shown for CODEX-measured and UPMC model-predicted cell types on the two validation datasets.

### Training on only one coverslip

Because highly multiplexed fluorescence imaging platforms like CODEX are more resource-intensive than standard fluorescence immunochemistry imaging, one might wish to image only one coverslip with CODEX, and then train a model to impute additional biomarkers on other coverslips imaged with a 7-plex system. We experiment with only training the imputation model on one coverslip (24% of the entire training data), and report that performance from this model retains the same cell type prediction performance (Supp. Table 1a). Even in this low data regime, the model can still robustly impute cell types without sacrificing performance.

### 7-UP leverages cell morphology

Finally, we verify that the deep learning model learns morphology features useful for imputing biomarkers for single cells beyond the mean expression values. Using only average cell expression values as input features, our method achieves an average Pearson correlation coefficient (PCC) of 0.474 across all predicted biomarkers in the UPMC-HNC dataset. When adding additional morphology features, the performance improves to 0.534 PCC (Table 1). Similarly, when determining cell types, a model which uses only the average expressions of seven biomarkers achieves an average patchwise weighted F1 score of 0.667. In contrast, the model with biomarkers imputed using morphology features achieves an F1 score of 0.727.

Additionally, to demonstrate the usefulness of the context channels used in the deep learning model, we performed an ablation experiment (Supp. Table 1b) where we evaluated a model trained using the context channels, and a model trained without (only using the single-cell image). We observe an improvement (0.016 PCC and 0.014 F1) with the inclusion of context channels, indicating that features from the cell’s neighborhood are useful in determining information about the cell.

As an example, classification performance on vessel cells increases from 0.44 to 0.67 F1 score when including morphological features. Of the cells incorrectly classified without morphology, 70% were predicted as stromal cells. Supp. Figure 2 visualizes examples of cells that were corrected with the inclusion of morphology. Though vessel cells and stromal cells share a similar protein expression composition (Vimentin, aSMA, CollagenIV, CD47), vessel cells uniquely express CD31 and CD34. Indeed, the model with morphology more accurately predicts CD31 (PCC: 0.561 vs 0.416) and CD34 (PCC: 0.586 vs 0.429). We can infer that the model was able to better predict the expression of these two biomarkers with morphological information of the 7 biomarkers than with only average expression.

## Discussion

High-plex immunofluorescence techniques like CODEX enable an unprecedented understanding of TME and tissue architecture but have seen limited clinical (diagnostic or prognostic) utility due to their cost and data generation times. On the other hand, standard IF or immunostaining workflows, which image between 1 to 7 biomarkers, are widely available. 7-plex mIF panels are becoming more common in clinical settings. Our proposed method aims to unlock the richer TME representations available with CODEX by up-leveling existing 7-plex data through learning biomarker co-expression and morphological patterns.

*7-UP* demonstrates that a small subset of biomarkers can contain sufficient signal to reconstruct a much larger subset of biomarkers. For instance, some biomarkers regularly co-express with other biomarkers (e.g., CD20 and CD21 in B cells), while others can be inferred from the cell’s morphology (e.g., the nucleus and cytokeratin expression of a proliferating tumor cell may indicate Ki67 expression). Indeed, our results suggest that learning these relationships is useful and that the imputed biomarker expressions are reliable enough to be used in place of CODEX-measured expressions for the primary tasks of resolving cell types and predicting phenotypic outcomes.

The panel selection procedure in Figure 1a demonstrates one method for selecting input biomarkers, which does so by maximizing the average reconstruction accuracy across all other CODEX-measured biomarkers. In scenarios where multiplex imaging data has been previously imaged and collected, *7-UP* can be deployed directly on the pre-defined subset of biomarkers, thus removing the need for panel selection.

The ability to determine a subset of biomarkers *in silico* 1) gives users immediate access to a larger set of biomarkers beyond what has been experimentally measured, and 2) frees up resources to measure more novel and biologically relevant biomarkers. Thus, in addition to up-leveling 7-plex systems, *7-UP* can also push CODEX systems from ~40 biomarker measurements to 60 or more, enabling even greater cell type differentiation and disease characterization.

### Limitations

While some biomarkers are imputable with a high degree of confidence, others are not as easily predicted. This is a consequence of the inherent limitations of a 7-plex panel. Intuitively, increasing the panel beyond seven biomarkers would increase the number of strongly predicted biomarkers, but would also surpass the technical limitation of most clinical multiplex workstations. Additionally, since biomarkers are differentially expressed based on their unique TME, training on a variety of disease contexts is key to ensuring generalizability. Picking an informative panel of biomarkers is also an important decision, and ought to reflect the nature of the disease and TME that one wishes to understand.

## Methods

### CODEX data collection

All samples are prepared, stained, and acquired following CODEX User Manual Rev C (https://www.akoyabio.com).

#### Coverslip preparation

Coverslips are coated with 0.1% poly-L-lysine solution to enhance adherence of tissue sections prior to mounting. The prepared coverslips are washed and stored according to the guidelines in the CODEX User Manual.

#### Tissue sectioning

formaldehyde-fixed paraffin-embedded (FFPE) samples are sectioned at a thickness of 3-5 μm on the poly-L-lysine coated glass coverslips.

#### Antibody conjugation

Custom conjugated antibodies are prepared using the CODEX Conjugation Kit, which includes the following steps: (1) the antibody is partially reduced to expose thiol ends of the antibody heavy chains; (2) the reduced antibody is conjugated with a CODEX barcode; (3) the conjugated antibody is purified; (4) Antibody Storage Solution is added for antibody stabilization for long term storage. Post-conjugated antibodies are validated by SDS-polyacrylamide gel electrophoresis (SDS-PAGE) and quality control (QC) tissue testing, where immunofluorescence images are stained and acquired following standard CODEX protocols, then evaluated by immunologists.

#### Staining

CODEX multiplexed immunofluorescence imaging was performed on FFPE patient biopsies using the Akoya Biosciences PhenoCycler platform (also known as CODEX). 5 μm thick sections were mounted onto poly-L-lysine-treated glass coverslips as tumor microarrays. Samples were pre-treated by heating on a 55 °C hot plate for 25 minutes and cooled for 5 minutes. Each coverslip was hydrated using an ethanol series: two washes in HistoChoice Clearing Agent, two in 100% ethanol, one wash each in 90%, 70%, 50%, and 30% ethanol solutions, and two washes in deionized water (ddH2O). Next, antigen retrieval was performed by immersing coverslips in Tris-EDTA pH 9.0 and incubating them in a pressure cooker for 20 minutes on the High setting, followed by 7 minutes to cool. Coverslips were washed twice for two minutes each in ddH2O, then washed in Hydration Buffer (Akoya Biosciences) twice for two minutes each. Next, coverslips were equilibrated in Staining Buffer (Akoya Biosciences) for 30 minutes. The conjugated antibody cocktail solution in Staining Buffer was added to coverslips in a humidity chamber and incubated for 3 hours at room temperature or 16 hours at 4 °C. After incubation, the sample coverslips are washed and fixed following the CODEX User Manual.

#### Data acquisition

Sample coverslips are mounted on a microscope stage. Images are acquired using a Keyence microscope that is configured to the PhenoCycler Instrument at a 20X objective. All of the sample collections were approved by institutional review boards.

#### Datasets

The UPMC-HNC and Stanford-HNC datasets have one held-out coverslip for model validation and one held-out coverslip for model evaluation. The Stanford-CRC dataset has half of one coverslip randomly split and held out for model validation and one held out for model evaluation.

### Choice of Input Biomarkers

Our 7-UP framework can be applied to any set of input biomarkers, though the imputation performance improves if the input markers are particularly informative. Concrete autoencoder^22^ is an unsupervised neural network that determines the subset of biomarkers that are most useful for reconstructing the entire CODEX panel (Figure 1a). The Concrete autoencoder takes a full set of input biomarker expressions and outputs a feature importance score for each biomarker. This approach achieves very similar results when compared to a naive greedy algorithm (iteratively including the most important biomarkers in the model), but is more computationally efficient.

### Biomarker expression preprocessing

Single cell expression was computed for each biomarker by 1. applying a deep learning cell segmentation algorithm (DeepCell)^27^ on the DAPI biomarker channel (nuclear stain) to obtain nuclear segmentation masks; 2. successively dilating segmentation masks by flipping pixels each time with a probability equal to the fraction of positive neighboring pixels (repeated 9 times); 3. computing the mean expression value across pixels within the single cell; and 4. normalizing the expression values across all cells in a sample using quantile normalization and arcsinh transformation followed by a z-score normalization:

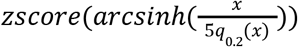

Where *zscore* is defined given μ and σ, the mean and standard deviation across all cell expression values in the sample:

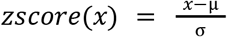

*x* is the vector of a biomarker’s values in a sample, *arcsinh* is the inverse hyperbolic sine function; and *q*_0.2_(*x*) is the 20th percentile of *x*.

### Image patch generation

After preprocessing (tile & cycle alignment, stitching, deconvolution, and background correction) CODEX data is available as multichannel OME-TIFF files, with each image channel corresponding to the fluorescence signal (expression) of a distinct biomarker probe. To prepare the input image patches for the deep learning model, we perform the following: All pixel values for a biomarker in a sample are normalized using ImageJ’s AutoAdjust function^28^. An image patch (224px by 224px) is then generated for each cell, for each biomarker, in the region. Each patch consists of three channels. The first channel contains the segmented cell only, rescaled 3x, centered with zero padding around the cell; the second channel contains a crop of the neighborhood (~20 cells) centered around the cell (~84 μm) at 1x scaling, and the third channel contains a crop of the neighborhood (~80 cells) at 0.5x scaling also centered around the cell (~42 μm). These channels are visualized in Figure 1b. As a reference, all coverslips are imaged at a resolution of 0.3775 μm per pixel.

### Deep learning model

We trained a ResNet-50^24^ deep learning model to learn cell shape features as well as spatial information of cell neighborhoods. We find that training the model on cell type classification enables it to learn an effective morphology featurizer. We start with a model with weights pre-trained on the ImageNet dataset^29^. The model takes as input a 224×224×21 size tensor, where the 21 channels correspond to stacking 7 input biomarkers with 3 feature channels each. The last layer is modified to classify over one-hot encoded cell types. The model is trained with categorical cross-entropy loss, and a cell-wise F1 score is computed at each validation step. The learning rate is initialized at 1e-4, and decays by a factor of 0.2 if the validation F1 score does not improve over 5000 steps. Training stops after 75k steps of no improvement, and the model with the highest validation F1 score is chosen. To improve model robustness, we trained an ensemble of five identical models with different random weight initializations and computed the mean prediction across all models to obtain a final model score. All models were implemented and trained using Pytorch^30^, a Python deep learning framework.

### Biomarker imputation model

XGBoost^25^, a gradient boosting decision tree algorithm shown to achieve top-performance in tabular data regression, is used for imputing single-cell biomarker expressions. The model takes as input the cell expression values of the seven input biomarkers, along with, in the case of adding morphology information, a probability vector corresponding to cell type predictions from the deep learning model. It is then trained to jointly predict the expression values of the remaining biomarkers. We find that directly using the output probabilities improves model performance more than using the final featurization layer. We used squared error loss, a learning rate of 0.1, 500 estimators, a max depth of 3, a per-tree column sampling of 0.7, and GPU accelerated training. All other hyperparameters are default settings in the XGBoost Python library.

### Dimensionality reduction

To visualize the concordance of CODEX-measured and imputed biomarkers, we randomly sample 10,000 cells with CODEX-measured biomarker values and 10,000 cells with imputed biomarker values and fit a UMAP^31^ dimensionality reduction model on the combined set. We then plot the projected 2D data points separately and color them by ground truth cell types, expression of CD45, and expression of CD20 (the latter two use the 75th percentile expression value as a binary threshold). The UMAP model is trained using a GPU-accelerated implementation^32^ with default settings.

### Patchwise Metrics

Given the naturally high degree of intercellular expression variation within local neighborhoods of cells, we report biomarker and cell type predictions aggregated within a local cell neighborhood. Supp. Figure 2 shows the relationship between the choice of cell neighborhood patch size (in pixels) and the average PCC and F1 score. The patchwise PCC of a biomarker is computed as the Pearson correlation coefficient between the CODEX-measured and imputed patchwise average expressions. Patchwise F1 is computed by considering a patch as positive if at least one cell is assigned to that cell type, and then calculating the F1 score across patches.

### Cell type ground truth and predictions

To produce cell type labels, we first obtained a cells-by-features biomarker expression matrix - for each marker, we took the average signal across all pixels in a segmented cell. This matrix was normalized and scaled as described above, then principal component (PC) analysis was performed. We constructed a nearest-neighbor graph (k = 30) of cell expression in PC space with the top 20 PCs, then performed self-supervised graph clustering^33^ on the result. Clusters were manually annotated according to their cell biomarker expression patterns. This procedure was performed on a subset of 10,000 cells and subsequently used to train a kNN algorithm. This algorithm was used to transfer labels to the entire dataset.

In our experiments where we generate cell type predictions based on the imputed biomarkers, we use this trained kNN algorithm and substitute the subset of expressions for which we are imputing with the imputed values from the ML model. Thus, for the UPMC-HNC dataset with 41 total biomarkers, 7 biomarkers will be the CODEX-measured values, and 33 biomarkers will be imputed.

### Survival outcome prediction

Additionally, three phenotypic patient outcomes from the UPMC dataset are evaluated: survival status (No Evidence of Disease (NED) versus Died of Disease (DOD)), HPV status (a significant indicator of cancer prognosis), and recurrence (if the cancer recurs within 5 years after diagnosis).

We used a graph neural network (GNN)-based model^26^ trained on using cell types to predict patient phenotypic outcomes. This model transforms the structure of each sample into a graph network, where cells are connected by edges to neighboring cells. It then pools information about the neighboring cells’ cell types to output an outcome probability score for each cell. The sample predictions are generated by averaging the scores across all cells in that sample. We evaluated models that have been trained on three patient phenotypic outcomes: survival status, HPV status, and recurrence. To validate the utility of our imputed cell types, we replace the original annotated cell type labels with the predicted cell types produced from the imputed biomarkers. The results are reported in Figure 4, where we see that performance on these three tasks is comparable between using the imputed biomarkers and the CODEX-generated biomarkers.

## Acknowledgments

J.Z. is supported by NSF CAREER 1942926. K.S. is supported by a Knight-Hennessy Fellowship.

## Data and Code Availability

All code used to produce the results in this paper is available at https://gitlab.com/enable-medicine-public/7-up. Data will be available upon request.

## Supplemental Tables and Figures

**Supplemental Table 1:**
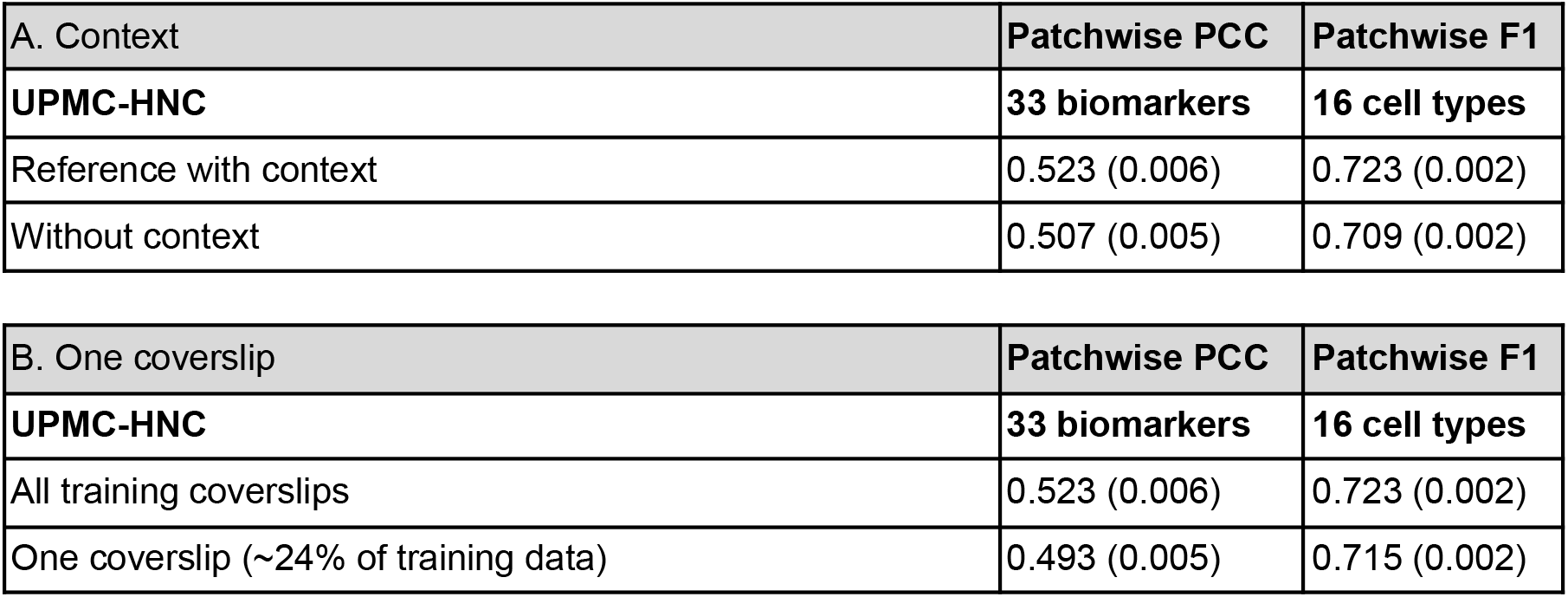
7-UP ablation studies. In Table A, the 7-UP model is trained with only the “Cell only” channel as described in Figure 1b, thus discarding the two context channels. The performance of this model is reported in the second row (“Without context”), and compared to a model trained with all three channels (“Reference with context”). In Table B, the 7-UP model is trained using only data coming from one coverslip, which represents about a quarter of the total training data. In both experiments, the ablated model performs comparatively similar to the reference model.

| A. Context | Patchwise PCC | Patchwise F1 |
| --- | --- | --- |
| **UPMC-HNC** | **33 biomarkers** | **16 cell types** |
| Reference with context | 0.523 (0.006) | 0.723 (0.002) |
| Without context | 0.507 (0.005) | 0.709 (0.002) |

**Supplemental Table 1:** 7-UP ablation studies.
| B. One coverslip | Patchwise PCC | Patchwise F1 |
| --- | --- | --- |
| **UPMC-HNC** | **33 biomarkers** | **16 cell types** |
| All training coverslips | 0.523 (0.006) | 0.723 (0.002) |
| One coverslip (~24% of training data) | 0.493 (0.005) | 0.715 (0.002) |

**Supplemental Table 2:**
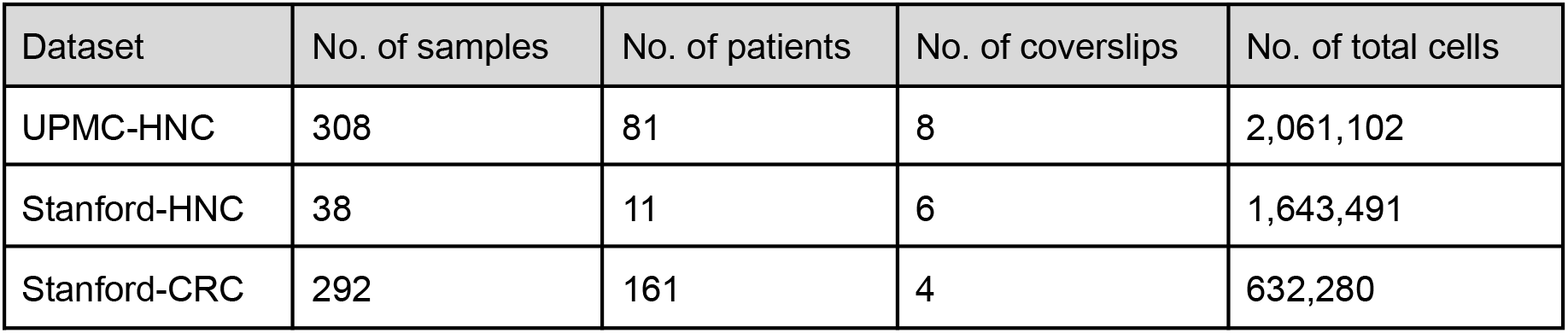
Descriptions of the three CODEX datasets used for training and evaluating 7-UP.

**Supp. Figure 1:**
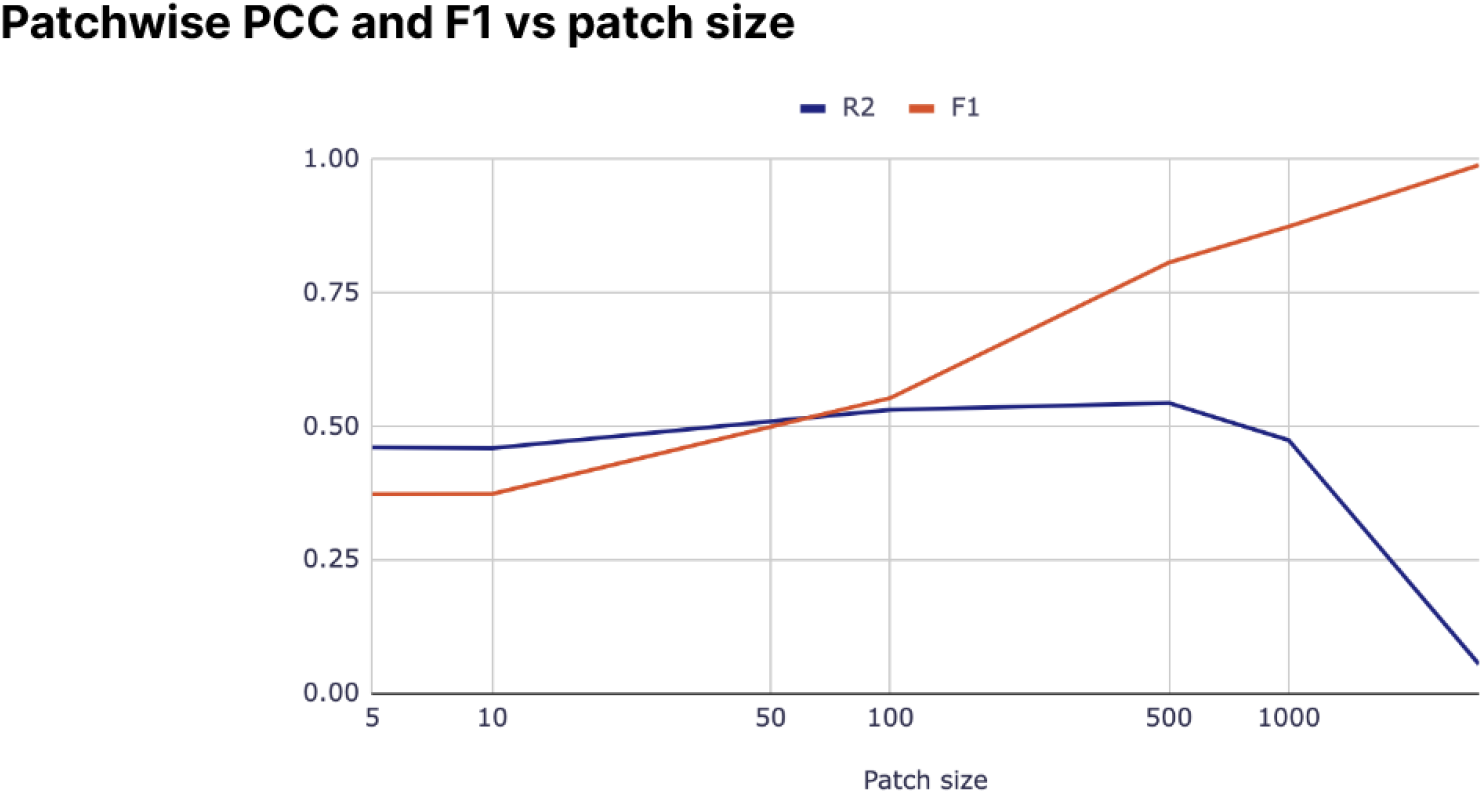
The effect of patch size (in pixels) on average patchwise PCC and F1. We use a patch size of 100 pixels when computing patchwise metrics.

**Supp. Figure 2:**
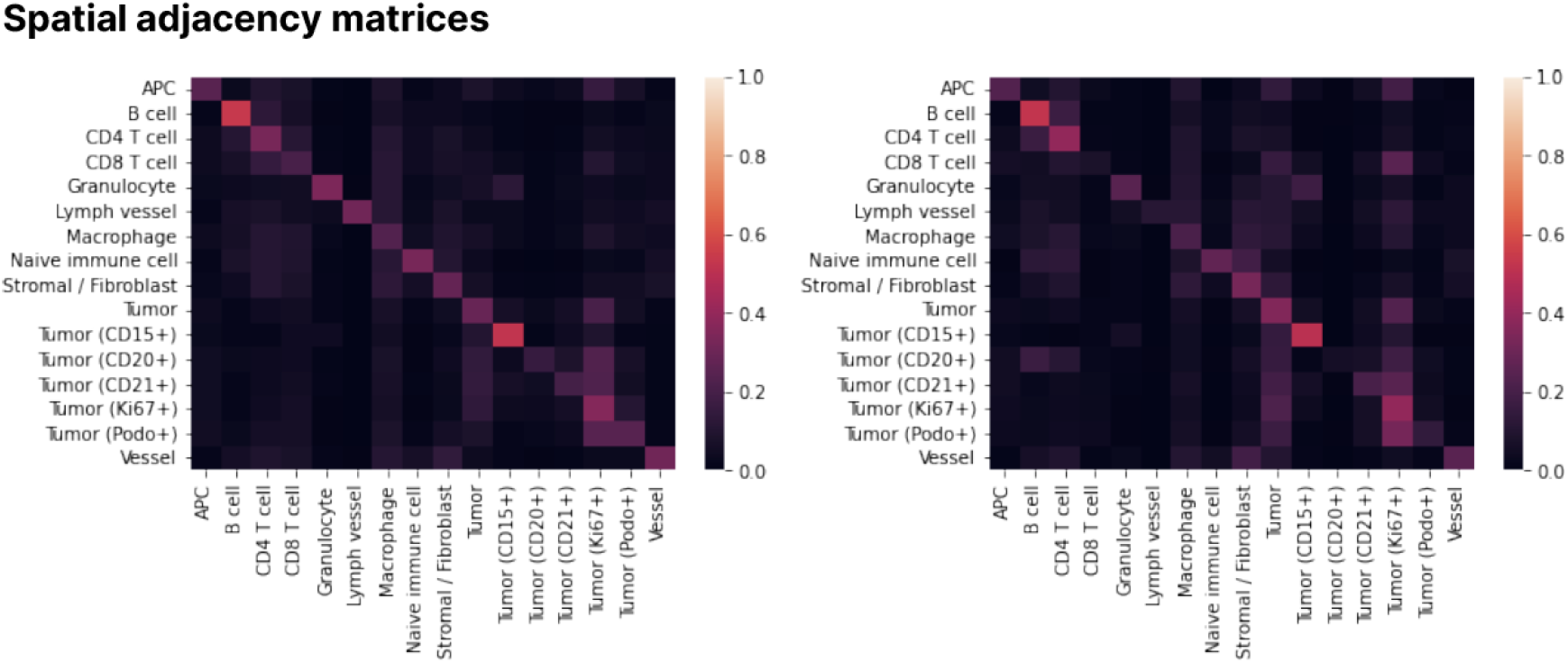
Spatial adjacency matrix agreement. To assess how well the cell type predictions preserve local neighborhood structures, we consider spatial adjacency matrices, which show the row-normalized counts of cell-cell neighbors within the UPMC test set.

**Supp. Figure 3:**
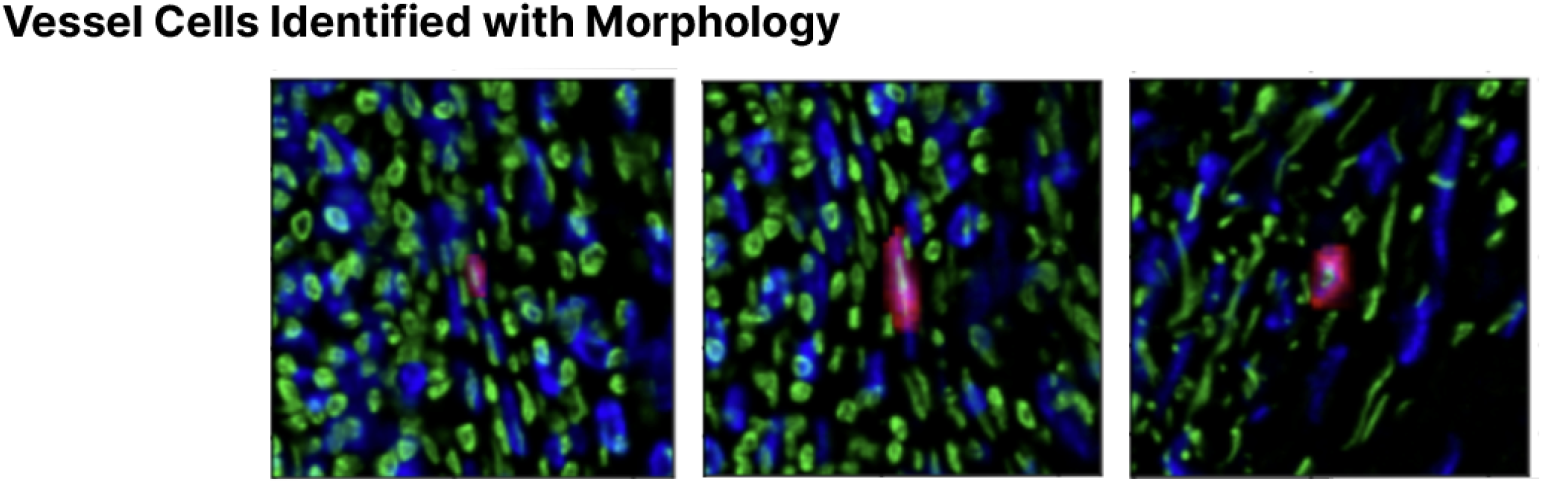
Vessel cells identified with morphology. Three patches of vessel cells were incorrectly classified as stromal cells but correctly classified with the inclusion of spatial information. In each patch, the DAPI stain is shown in three spatial scales: the cell morphology is presented in red, the 1x resolution context around the cell is shown in blue, and the 0.5x resolution context around the cell is shown in green.

**Supp. Figure 4:**
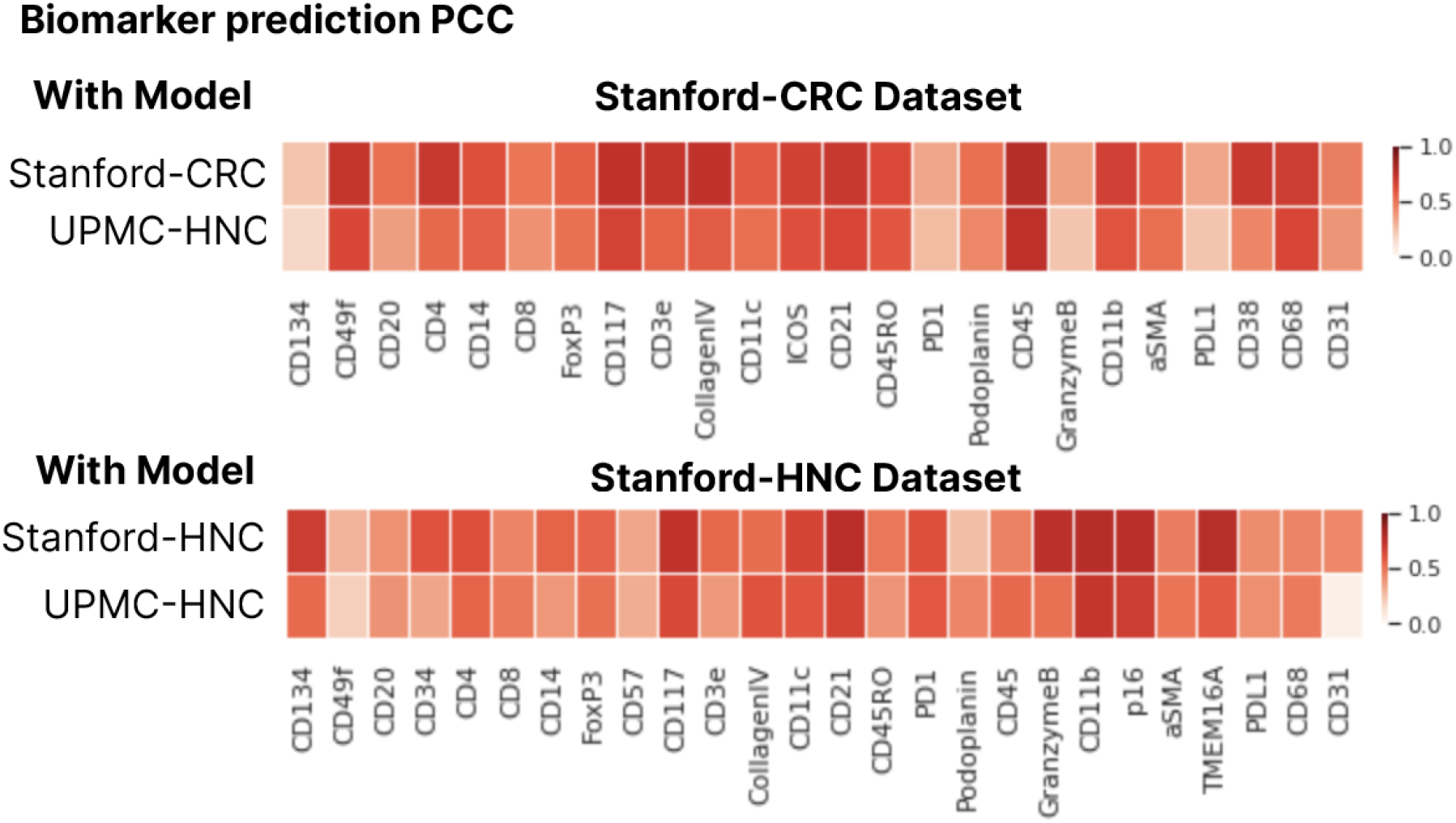
A breakdown of patchwise PCC per biomarker is visualized for each cross-site evaluation.

